# Robust foreground detection in somatic copy number data

**DOI:** 10.1101/847681

**Authors:** Aditya Deshpande, Trent Walradt, Ya Hu, Amnon Koren, Marcin Imielinski

**Affiliations:** Tri Institutional PhD Program in Computational Biology and Medicine, Graduate School of Medical Sciences, Weill Cornell Medicine, 1300 York Avenue, New York, NY 10065, USA; Weill Cornell Medical College, 445 East 69th Street, New York, NY 10065, USA; Department of Molecular Biology and Genetics, Cornell University, Ithaca, NY 14853, USA; Pathology and Laboratory Medicine, Weill Cornell Medicine, 1300 York Avenue, New York, NY 10065, USA; New York Genome Center, 101 Avenue of the Americas, New York, NY 10013, USA; Institute for Computational Biomedicine, Weill Cornell Medicine, 1300 York Avenue, New York, NY 10065, USA

## Abstract

Sensitive detection of somatic copy number alterations (SCNA) in cancer genomes is confounded by “waviness” in read depth data. We present `dryclean`, a signal processing algorithm to optimize SCNA detection in whole genome (WGS) and targeted sequencing platforms through foreground detection and background subtraction of read depth data. Application of `dryclean` to WGS demonstrates that WGS waviness is driven by replication timing. Re-analysis of thousands of tumor profiles reveals that `dryclean` provides superior detection of biologically relevant SCNAs relative to state-of-the-art algorithms. Applied to *in silico* tumor dilutions, `dryclean` improves the sensitivity of relapse detection 10-fold relative to current standards. `dryclean` is available as an R package in the GitHub repository https://github.com/mskilab/dryclean

## Background

Accurate somatic copy number alteration (SCNA) detection is a crucial aspect of precision oncology and cancer genome analysis. Therapeutically actionable SCNAs include trastuzumab-sensitizing *ERBB2* amplifications in breast cancer (1) and *CDKN2A* deletions or *CDK4* amplifications identifying responders to *CDK4/6* inhibitors in glioblastoma (2). SCNAs resulting in the loss of *RB* and *TP53* are associated with later “small-cell conversion”, a mechanism of aggressive erlotinib-resistant relapse in lung adenocarcinoma (3). In a modern oncology practice, the decision to provide or deny a certain therapy may be heavily influenced by sequencing results (4). Consequently, it is vital that these tests achieve an adequate level of sensitivity. Accurate SCNA detection is also crucial for enabling discovery of recurrent patterns of structural variation or genotyping SCNAs in samples where somatic clones are highly diluted against a euploid background (e.g. solid and liquid biopsy relapse monitoring, clonal hematopoiesis detection) (5–7).

A key challenge in SCNA inference from DNA sequencing experiments is noise or bias in read depth data (8). SCNA inference relies on an affine relationship between copy number (CN) and read depth to identify segments of discrete CN states. This relationship is distorted by assay– and batch-specific effects, including mappability bias, GC bias, library quality, input quantity, and heterogeneity in capture probe affinity (9, 10). Though many methods exist for SCNA inference in whole genome (WGS), whole exome (WES), or targeted panel (TP) sequencing, most take the tumor-normal ratio (TNR) as input (6, 11–16). The reason for this normalization, in theory, is to remove systematic biases since both tumor and normal samples are processed similarly (e.g. similar capture probes, library preparation, read lengths).

As a ratio of two noisy signals, however, the TNR is sensitive to subtle or stochastic differences in batch effects, including low-quality normal libraries or specimens (11). Furthermore, the presence of inherited CN events can lead to puzzling patterns in the TNR, such as foci of apparent CN gain when a region containing a focal germline deletion undergoes loss of heterozygosity. Consequently, reliance on TNR may lead to misinterpretation of data (false negatives and false positives). Finally, a matched normal may be absent for certain samples, such as a cancer cell line or archival formalin-fixed paraffin-embedded (FFPE) tissue. Here, computation of TNR from an arbitrary or composite diploid sample may provide inadequate normalization or even add distortion to the read depth signal.

An ideal read depth normalization method would flatten assay– and batch-specific biases (termed “background”) while retaining transitions associated with CN changes (termed “foreground”). Previous approaches for read depth normalization include supervised algorithms that explicitly correct for known factors, specifically GC bias and mappability (10, 17)). As we show, GC and mappability correction fails to remove significant biases that plague both whole exome and whole genome sequencing read depth data, causing artifacts in downstream SCNA analyses. To overcome the significant limitations of these state-of-the-art pipelines, we have developed `dryclean`. `dryclean` learns a background model of biological and technical bias through the analysis of a panel of diploid normal samples (PON) using a signal processing method previously used for object detection in video data (18). Applying this model to tumor data, `dryclean` enriches for SCNA signal while subtracting the assay– and batch-specific background. Through reanalysis of large sequencing datasets from the Cancer Cell Line Encyclopedia (CCLE) and the Cancer Genome Atlas (TCGA), we reveal footprints of DNA replication in WGS background, improve the calling of biologically significant and clinically actionable SCNA, and demonstrate sensitive detection of SCNA in very dilute (< 1%) tumor samples (such as those that might be obtained during liquid or solid biopsy relapse monitoring).

## Results

### Robust foreground detection in read depth data

A central premise of `dryclean` is that genome-wide read depth data is shaped by an assay, batch, and sample specific background that arises from the combination of a small set of (unknown) factors, each affecting a large fraction of samples. In contrast, each tumor sample harbors a unique SCNA profile, comprising both broad (e.g. arm level gains and losses) and focal (e.g. double minute amplifications) variants that are rarely seen in normal samples.

To apply this intuition to cancer genomic read depth data, we borrow a powerful tool from computer vision and signal processing called robust principal components analysis (rPCA). A key previous application of rPCA is object detection in video surveillance (18). Here, a set of video images comprise a slowly changing background occasionally perturbed by a fast moving object, which defines the foreground of a subset of pixels in a handful of frames. Through the solution of an optimization, rPCA can decompose the matrix of *n* video images, each with *m* pixels, as the sum of a low rank (slowly changing) background matrix and sparse foreground matrix.

Similarly, we apply rPCA to detect cancer SCNA as a sparse and high amplitude perturbation against a relatively constant background defined by *n* – 1 normal diploid samples, each comprising read depth data across a set of *m* genomic bins. Formally, we decompose the input matrix 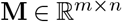 of read depth data across *n* – 1 normal samples and one tumor sample into a low-rank matrix 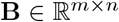 and sparse matrix 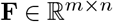 through the solution of a constrained optimization:

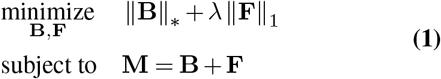

where λ > 0 is a regularization parameter. For a matrix 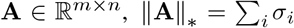 is the nuclear norm and 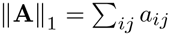 is the *L*_1_ norm of a vectorized **A** matrix. Here, *σ_i_* is the *i*th singular value of matrix **A** and *a_ij_* is the *ij* component of matrix **A**. In practice, we separate the solution of Eq. 1 into a computationally intensive “training” step and faster “inference” step (Figure 1, see Methods). The training step directly solves Eq. 2 for a matrix 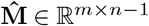 of *n* – 1 diploid samples to obtain 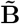 and 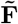. The inference step computes the projection **b** of a new tumor sample 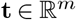 on the low-rank “background” column space of 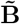 and a sparse residual vector **f** that we call “foreground” (Figure 1). As we show below, foreground vectors represent SCNA and background vectors represent assay– and batch-specific technical and biological biases.

**Fig. 1.**
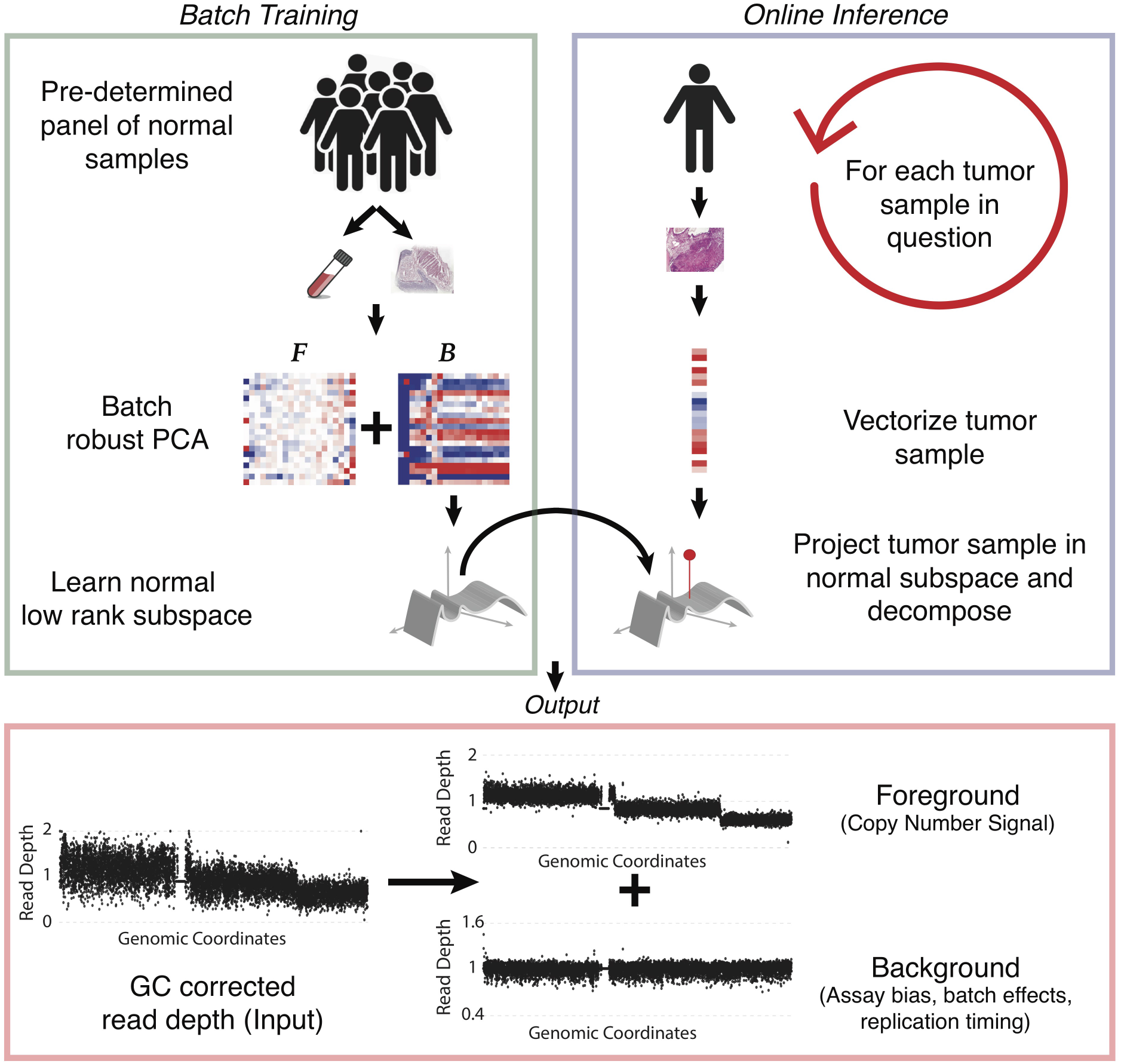
dryclean schematic. The `dryclean` pipeline separates foreground (SCNA signal) from background (technical and biological bias) in read depth data by adapting an algorithm used to detect objects in video surveillance data. `dryclean` learns a low rank subspace across a panel of normal (PON) samples using rPCA and applies this to a tumor sample to infer SCNA. The three boxes indicate stages of the `dryclean` pipeline. First, (green box) a PON is created from log transformed, zero centered, GC and mappability corrected read depths across a set of normal samples and genomic windows. Batch rPCA decomposition yields a low rank **B** matrix, whose columns span a subspace of biases that define the assay “background”. A second step (purple box) projects a tumor onto the background subspace to identify two vectors: the low-rank projection (**b**) representing background bias and sparse residual (**f**) corresponding to foreground or SCNA (red box).

### dryclean foreground enriches SCNA signal across sequencing platforms

To gain intuition into the decomposition performed by `dryclean`, we compared unsegmented WES and WGS read depth data for a TCGA tumor and normal pair (TCGA-02-2483). For this analysis, we compared the GC and mappability corrected read counts (17) for bins present in both WGS and WES. In addition, we superimposed copy number signal from the segmented SNP6 gold standard SCNA data on to these bins. Comparison of WES and WGS TNR demonstrated poor inter-platform correlation (*R*^2^ = 0.094, Figure 2A) and poor visual evidence of discrete CN states in either dataset. This was further corroborated by a weak separation of the regions with different segmented CN states (as ascertained by SNP6). This correlation and CN separation was similarly poor when we examined the tumor-only GC and mappability corrected read counts, which comprise the numerator to the TNR and the input to `dryclean` (i.e. the **m** vector) (*R*^2^ = 0.05, 2B).

**Fig. 2.**
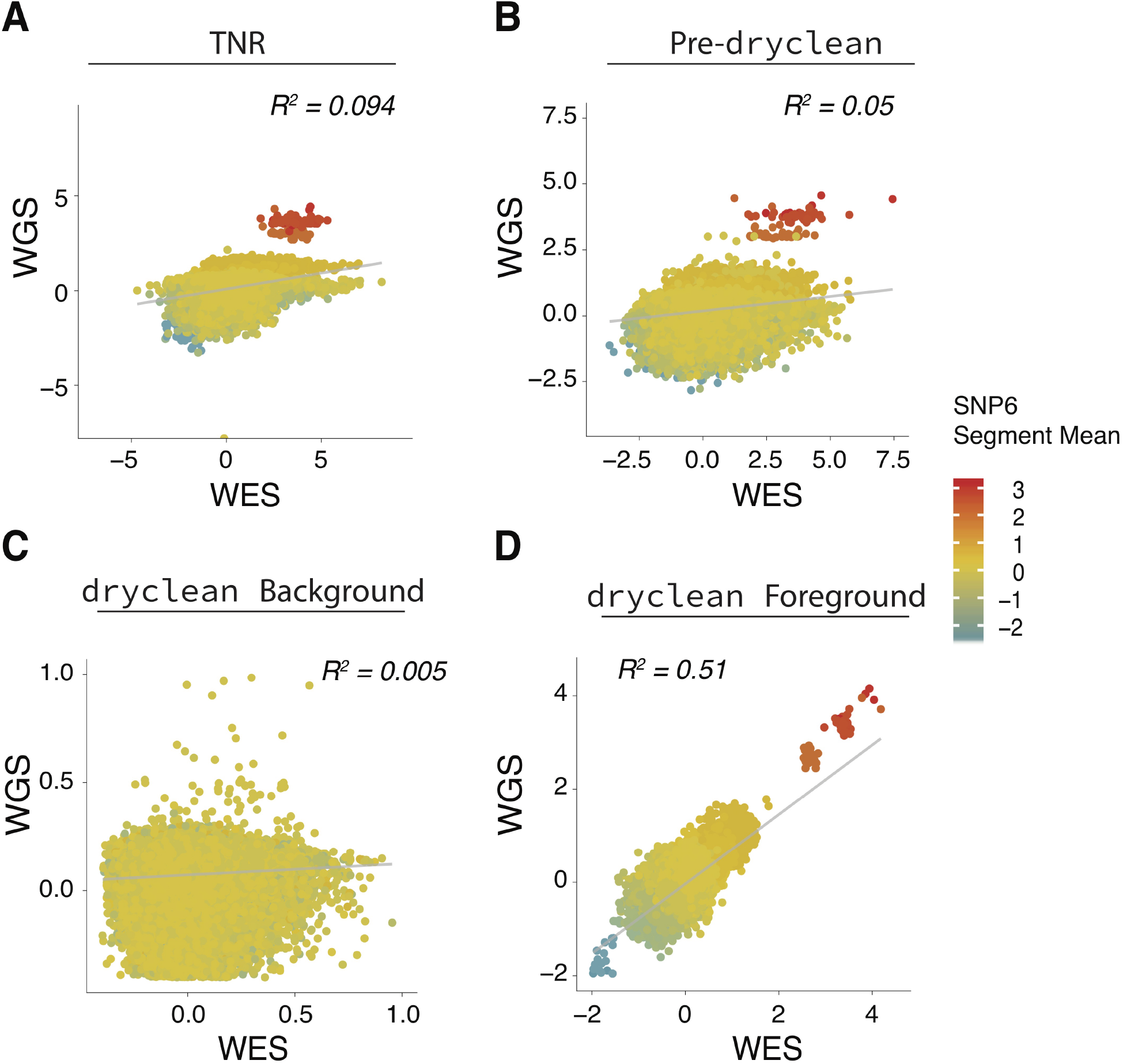
dryclean foreground reveals SCNA signal across diverse sequencing assays. For (A-D), WES and WGS read depth data are shown for a TCGA tumor and normal pair (TCGA-02-2483). Each point on the scatter plot represent GC and mappability corrected (17) read count for bins present in both WGS and WES. The color of each point represents the copy number signal from segmented SNP6 gold standard data (log). (A) The TNR shows poor inter-platform correlation (*R*^2^ = 0.094) between WES and WGS, which suffer from different assay and batch specific biases. The copy number separation is poor and the low copy number regions are indistinguishable. (B) This correlation is similarly low even when we compare the pre-`dryclean`ed vectors representing tumor read depth following GC and mappability correction (*R*^2^ = 0.05). As with TNR, the copy states are weakly separated. (C) The `dryclean` inferred background for WES and WGS data (decomposing **m** data against their respective PON) represents assay and batch specific biases, which (as expected) demonstrates little correlation (*R*^2^ = 0.005) between platforms. Regions from all copy state are intermixed as we would expect from the background. (D) The `dryclean` inferred foreground shows improved correlation between WGS and WES. (*R*^2^ = 0.51) and captures discrete clusters of read depth representing SCNAs. Notable are the clearly distinct copy states that track well with SNP6 data.

We posited that `dryclean`-inferred background vectors for WES and WGS data represents distinct assay– and batch-specific biases. Confirming our hypothesis, the **b** vectors demonstrated little correlation (*R*^2^ = 0.005) between the two platforms and were intermixed with respect to copy state (Figure 2C). In contrast, the foreground `dryclean` solution showed improved correlation between WGS and WES (*R*^2^ = 0.51, Figure 2D) and a clean and visually discrete pattern of points consistent with CN signal from SNP6 data. These results suggest that the genomic background learned by `dryclean` can help reveal CN signal in high-density read depth data that is otherwise obscured by assay– and batch-specific bias.

### WGS waves reflect DNA replication

Though WGS theoretically comprises uniformly sampled fragments from a DNA extract, cancer WGS data often contain read depth oscillations of low frequency (≥ 100kbp) and varying amplitude, also termed “waves”. These waves, also observed in microarrays (19, 20), cause false positive calls in downstream segmentation algorithms that detect SCNA as read depth change points. WGS waviness is most commonly attributed to technical factors (e.g. GC bias) during library preparation (10). Previous work in non-malignant cells have also linked fluctuations in WGS read depth to the ongoing replication of cells (21). A key challenge in detecting replication driven fluctuations is separating them from SCNAs, which are abundant in aneuploid cancer cells.

To examine the relationship between `dryclean` output, copy number, and replication timing we analyzed a particularly “wavy” WGS-profiled CCLE cell line (G16478.KE-39, stomach cancer cell line (22)). We compared read depth signal before and after `dryclean` correction to an average of 6 lymphoblastoid cell line (LCL) replication timing profiles obtained from two 1KGP trios (21). Indeed, analysis of 1000 randomly selected genomic bins of the `dryclean` input showed non-negligible correlation (*R*^2^ = 0.16, Figure 3A) between replication timing z-scores and GC– / mappability-corrected read depth. This correlation was complicated by the presence of SCNAs, which defined discrete clusters of genomic bins (Figure 3A, colors represent CN from SNP6-derived segments). This replication timing correlation disappeared when comparing against the `dryclean`-processed read depth foreground (**f**) vector (*R*^2^ = 0.002, Figure 3B), where the colored data points separated primarily on the basis of copy number. In contrast, the `dryclean`-derived background (**b**) vector showed high correlation with replication timing (*R*^2^ = 0.45, Figure 3C) and absence of correlation with SNP6-derived copy number. The ability of `dryclean` to separate the GC-corrected input signal into an SCNA-enriched foreground and replication timing-enriched background was further demonstrated in Figure 3D, where the background vector tracked with LCL replication timing while the foreground change points co-located with concordantly oriented rearrangement junctions.

**Fig. 3.**
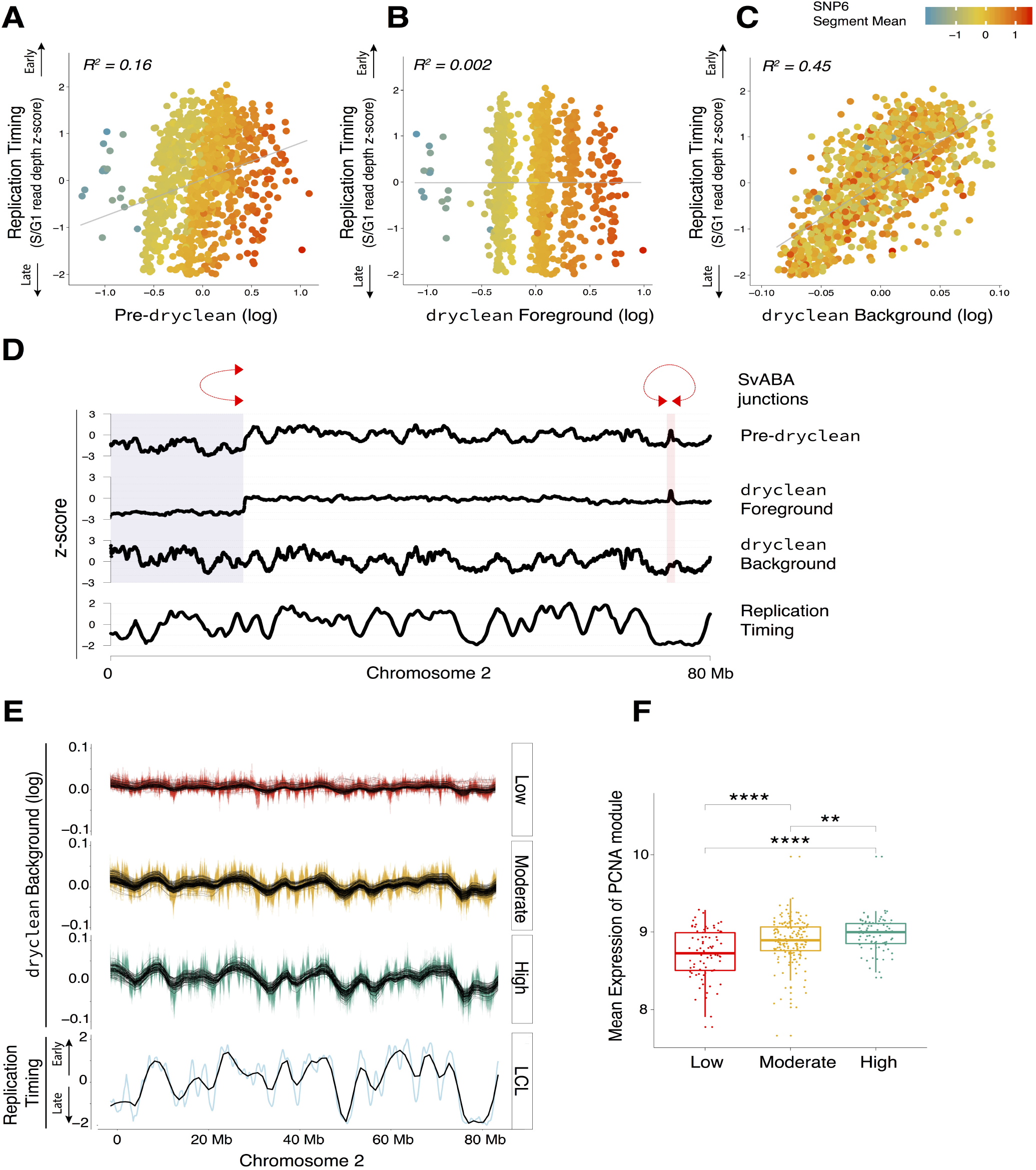
dryclean background reveals replication waves in cancer WGS. (A) Correlation of GC and mappability corrected log read counts i.e. pre-`dryclean` data at 1000 randomly sampled loci with LCL replication timing for a CCLE sample (G16478.KE-39, stomach cancer cell line) with high replication score (see text). Similar to Figure 2, the copy number signal from SNP6 data shows poor separation. (B) `dryclean` inferred foreground (**f** vector) correlates poorly with replication timing from LCL (*R*^2^ = 0.002), while show clear copy number states that track with SNP6. (C) `dryclean` inferred background (**b** vector) shows increased correlation (*R*^2^ = 0.45) with replication timing profile and poor separation of copy states. (D) Snapshot of these tracks across chromosome 2p demonstrates increased correlation of `dryclean` background with replication timing relative to input i.e Pre-`dryclean`, while `dryclean` foreground signal reveals SCNA’s, whose change points coincide with the locations of rearrangement junctions (red links above track). The blue shaded region highlights a deletion and red region highlights a focal amplification whose endpoints coincide with nearby concordant rearrangement junctions. (E) `dryclean`-inferred background vectors from 320 CCLE lines, grouped by quantiles into three tranches of low, medium, and high replication timing signal (see methods). Each colored track represent raw data. Black curves represent loess-smoothed fit for each sample. (F) Box plot of Proliferating Cell Nuclear Antigen (PCNA) gene expression module (described in text) demonstrates transcriptional programs associated with increased DNA synthesis in cell lines with high or medium vs low replication scores.

Expanding this analysis to a larger dataset, we observed variable correlation (*R*^2^ range from 0.2 to 0.6) between background **b** vectors computed for 320 CCLE cell lines (22) and the LCL replication profile reference (Supplementary Figure S1). To quantify the magnitude of replication-associated waviness, we computed a “replication score” as the covariance of **b** and LCL replication timing vectors and classified cell lines as high, moderate or low replicating. Figure 3E demonstrates variable degrees of `dryclean`-derived background **b** vector waviness and its correlation with LCL replication timing in an example region of chromosome 2p for high, moderate and low replicating CCLE lines.

To determine whether cell-line specific differences in replication-correlated waviness represent differences in S-phase fraction in cancer cell lines, we assessed the expression of the “meta-PCNA gene module” (23). This module comprises the top 1% of genes that are most positively correlated with the proliferating cell nuclear antigen (PCNA) expression across an independent set of 36 normal tissues (23). PCNA is an essential component of replication machinery and is a commonly used marker of proliferation. Figure 3F shows a significant increase in PCNA module expression between low, moderate, and high-replicating CCLE groups (Wilcoxon rank sum test p value = 1.9 × 10^−7^ between high and low, p value = 6.47 × 10^−5^ between moderate and low and p = 0.01933 between high and moderate).

### dryclean reduces bias and improves SCNA detection

We broadly evaluated `dryclean`’s ability to reduce bias in read depth data across a dataset of WGS, WES and simulated TP. Comparing two metrics of noise and waviness (DLRS, hypersegmentation, see Methods for details) across 1576 samples and two platforms (WES and TP), we saw significant and consistent reduction in both metrics for `dryclean`-processed read depth relative to TNR (Wilcoxon rank-sum test p value < 2.2 × 10^−16^ for all comparisons) (Supplementary Figure S2B-C). Moreover, for both WES and TP data, we identified a subset of TNR outliers that were “rescued” by `dryclean` (Supplementary Figure S2A). We observed similar reduction of noise metrics relative to TNR across 943 WGS cases (Supplementary Figure S2B-C). Figure 4A shows chromosome 13 of a bladder cancer (TCGA-DK-A1AA) WGS sample with very noisy coverage (DLRS= 0.145) and hypersegmentation that persisted despite GC bias and mappability correction. Following `dryclean` coverage correction, the DLRS value was lowered to 0.06 and SCNAs emerged visibly in the read depth data and subsequent segmentation.

**Fig. 4.**
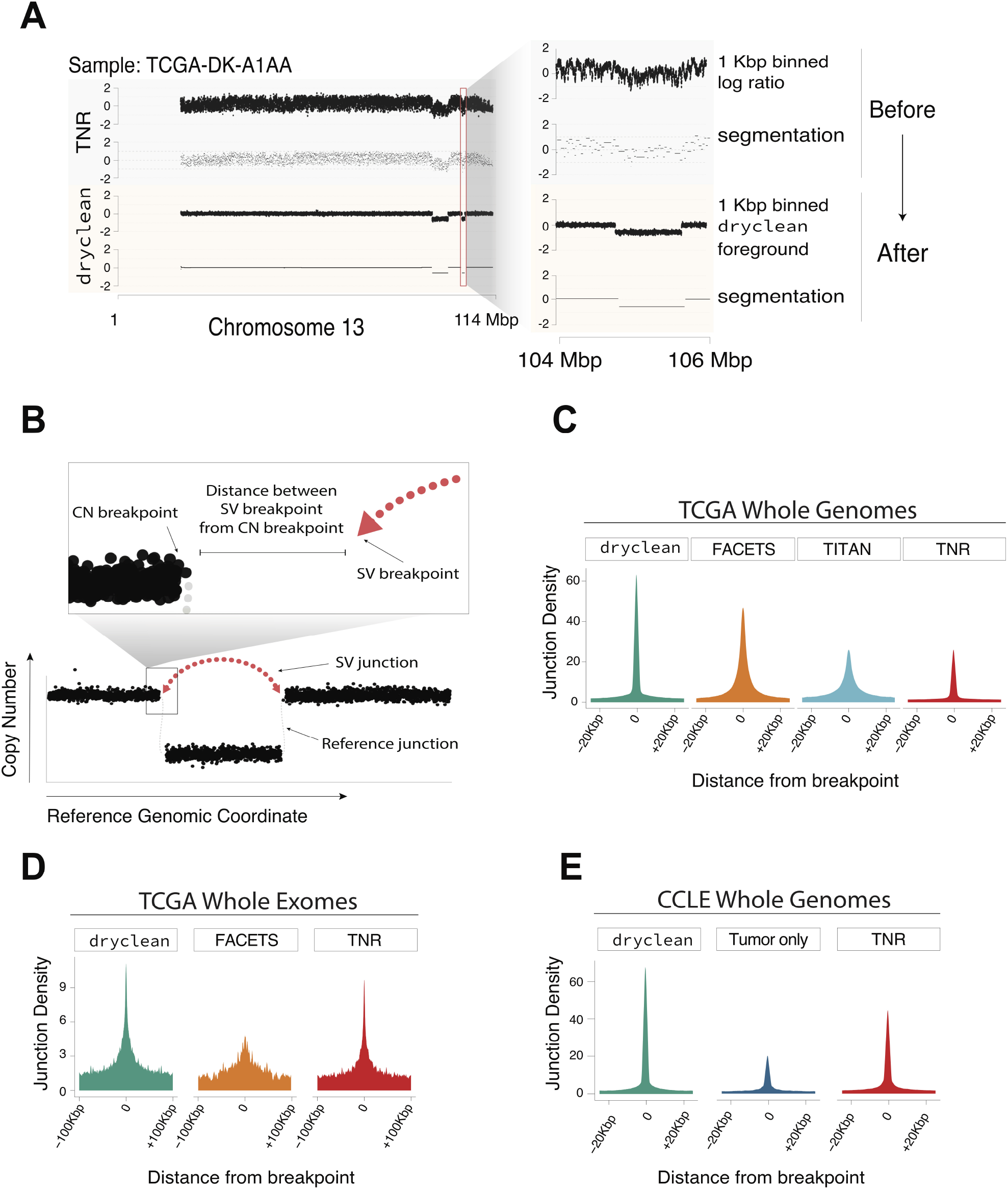
dryclean improves SCNA signal in WGS and WES read depth data. (A) Example of a TCGA WGS case with noisy / wavy TNR and hypersegmented CBS output. The `dryclean` inferred foreground demonstrates uniform coverage except for CN endpoints, yielding a dramatically improved segmentation. (B) Schematic of analyses in C-E, which rely on the premise that structural variant junction break points should occur at nearly all true SCNA. (C-E) Distribution of structural variation junctions around CN endpoints from TNR, FACETS, `dryclean` and TITAN in (C) TCGA WGS, (D) TCGA WES and (E) CCLE WGS data. CN end-points from `dryclean` segmentations have the highest and most focal density of junctions within 20 Kbp (WGS) and within 100 Kbp (WES) relative to other methods.

Since `dryclean` is not an SCNA caller *per se* but a method for reducing read depth bias, we combined it with a known segmentation method called circular binary segmentation (CBS, (24)). We compared the resulting SCNA calls (CBS-DC) against FACETS (14) or CBS combined with TNR (CBS-TNR), two representative state-of-the-art approaches. Using SNP6 segmentation as an orthogonal gold standard, we found that the locations of CBS-DC segment endpoints were significantly more concordant with SNP6 than CBS-TNR and FACETS results in both WES and TP data (Wilcoxon rank-sum test p value < 0.001 for all comparisons, Supplementary Figure S3A). In addition, the CBS-DC copy status (loss vs. neutral vs. gain, see Methods for details) at overlapping segments showed greater concordance with SNP6 than CBS-TNR and FACETS (Wilcoxon rank sum test p value < 2.2 × 10^−16^ for all comparisons) (Supplementary Figure S3B).

Since WGS can theoretically provide a more accurate estimate of CN than microarrays, SNP6 data does not provide an adequate “gold standard” for WGS-derived calls. Thus, to evaluate `dryclean`’s WGS performance, we used the presence of structural variant (SV) junctions (Figure 4B) as an orthogonal benchmark of SCNA accuracy.

Specifically, we quantified the distance between SvABA-derived (25) rearrangement junctions and SCNA segment end-points generated by `dryclean` vs. three state-of-the-art SCNA methods (CBS-TNR, FACETS, TITAN) (13, 14). Indeed, we found that `dryclean`-derived SCNA were focally enriched in SvABA junctions within a few Kbp of the segment end point (Figure 4C). We saw similar improvement in WES, where SV junctions were most highly enriched around `dryclean` breakpoints (Figure 4D) relative to other state-of-the-art approaches, including FACETS. Finally, application of `dryclean` to 320 CCLE cancer cell lines (22) lacking a matched normal demonstrated improved SCNA detection relative to two versions of CBS (FACETS and TITAN could not be compared, since they require a matched normal) (Figure 4E, see Methods). These results suggest that the read depth noise reduction achieved by `dryclean` enriches for signal associated with true SCNA across multiple sequencing platforms and sample types.

### dryclean provides superior detection of clinically actionable SCNA

We assessed `dryclean`’s ability to detect clinically-actionable SCNAs, i.e. those potentially informing prognosis and/or treatment either as standard of care or in the context of a clinical trial. We defined actionable events using annotations from Memorial Sloan Kettering Cancer Center’s OncoKB database (26) with the addition of some genes known to be prognostic in a particular cancer type (*CDKN2A* deletion (27) and *MDM2* amplification (28) in non-small cell lung cancer, *PTEN* deletion (29) in prostate adenocarcinoma). We computed precision and recall using the SNP6 annotation as the gold standard for both WES and TP analyses. As shown in Figure 5, F1 scores (harmonic mean of precision and recall) for `dryclean` matched or exceeded those for FACETS and TNR for several clinically relevant CNA for both WES and TP. Table S1 shows the sample counts used for the F1 calculations.

**Fig. 5.**
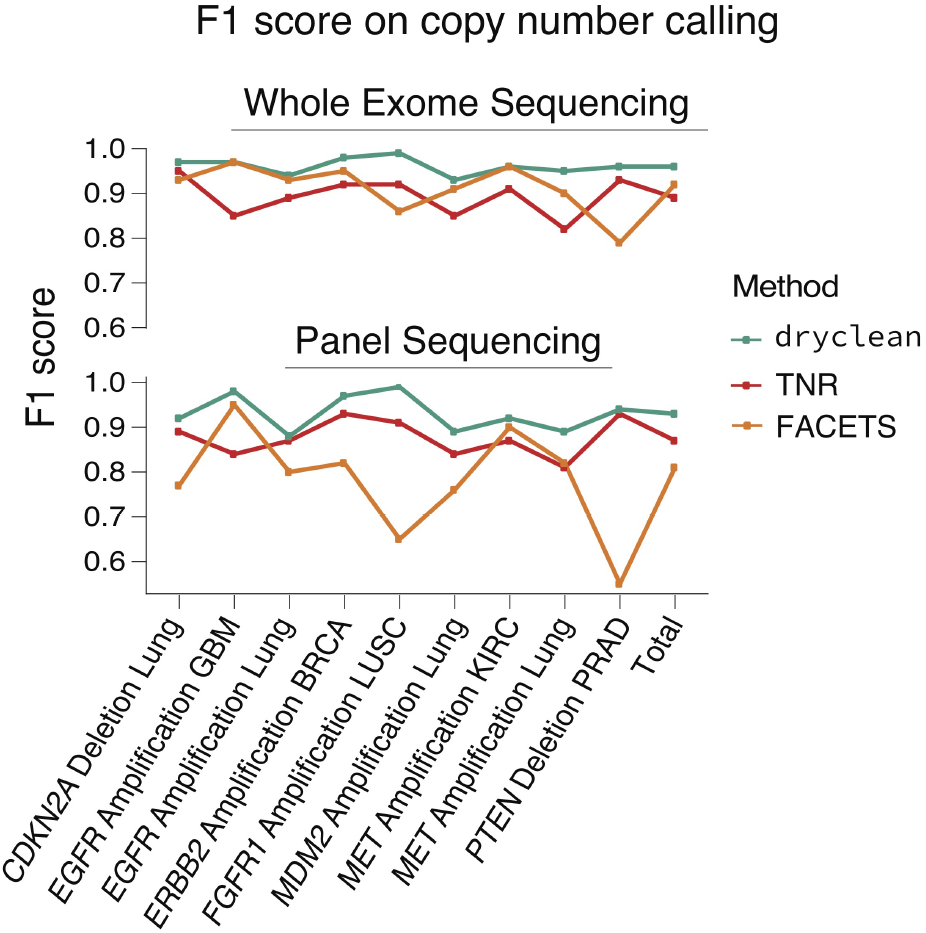
dryclean enables superior detection of clinically actionable SCNA. Upper panel shows F1 scores of different methods with respect to SNP6. `dryclean` shows the highest accuracy, with F1 scores exceeding 90% for all loci and cancer types. The lower panel shows the F1 score for TP. Lung includes lung adenocarcinoma and lung squamous cell cancers.

The presence of WES and TP bias obscured clinically important SCNA and resulted in important false negatives (Supplementary Figure S4A-C). This included focal *PTEN* loss and *MET* amplification, each of which was missed by TNR and FACETS. Supplementary Figure S4C provides a cautionary vignette on the use of matched normal samples in SCNA calling. Instead of calling a focal amplification in *EGFR*, which were seen in both `dryclean` and SNP6, both TNR and FACETS identified an artifactual deletion. Inspection of raw read counts in the matched normal sample (Supplementary Figure S5) showed that the “solid tissue normal” sample also had a broad and high amplitude *EGFR* amplification, possibly indicating a sample mix up or tumor-in-normal contamination. When this sample was used as a paired normal, it created a spurious drop in signal that was interpreted by downstream tools as a deletion. By modeling a broad PON, `dryclean` was immune to this artifact and agreed exactly with the SNP6 signal.

### dryclean detects SCNA in dilute tumor samples

The detection of somatic variants in circulating tumor DNA (ctDNA) enables non-invasive monitoring of relapse and resistance in patients receiving targeted drugs and/or chemotherapy (5, 30, 31). `dryclean`’s ability to enrich SCNA signal above an assay– and batch-specific background could be potentially leveraged to detect microscopic relapse or drug resistant driver genotypes in WGS of ctDNA or solid biopsies. SCNA-based relapse detection might be particularly sensitive in cases where a primary tumor SCNA profile was previously obtained with WGS, WES, or TP.

To explore this application for breast cancer relapse detection, we generated 30X tumor-normal *in silico* dilutions from five TCGA breast cancer cases with amplified *ERBB2* for which we had WGS, WES, and simulated TP data. We examined 5 dilution fractions (10%, 1%, 0.1%, 0.01% and 0%) in 10 replicates for changes in mean read depth at the *ERBB2* locus relative to pure germline DNA (0% mixture where the paired normal sample was diluted into a randomly selected blood sample).

Figure 6A depicts the signal from TNR and `dryclean`. We were able to detect significant signal in `dryclean` output at *ERBB2* for as low as 1 in 1000 tumor cell concentration (Wilcoxon rank sum test p value = 5.5 × 10^−4^). In contrast, TNR was unable to detect significant signal relative to negative control samples at lower than 1 in 100 tumor DNA fraction, below which changes in signal intensity were indistinguishable from noise. Receiver operating characteristic (ROC) curves (Figure 6B) demonstrated superior tumor detection with `dryclean` relative to TNR across various dilutions and sequencing assays.

**Fig. 6.**
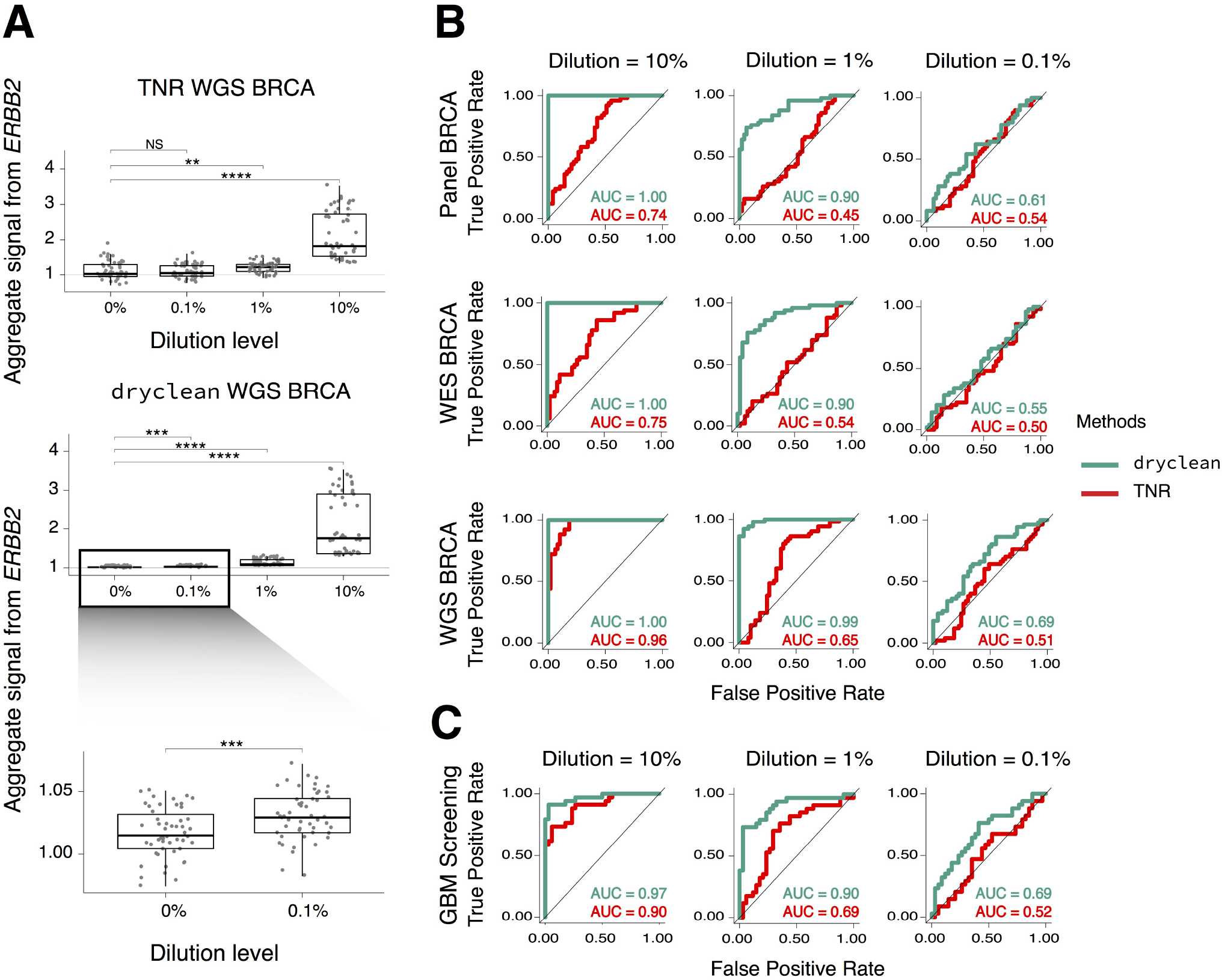
dryclean enables sensitive detection of dilute tumor DNA. (A) Aggregated intensity of `dryclean` foreground vs. TNR signal at the *ERBB2* locus is shown across various *in silico* dilutions of tumor in blood WGS. `dryclean` detects *ERBB2* amplification at as low as 0.1% tumor concentration (lower panel, zoomed), relative to control dilutions with 0 tumor DNA or results obtained from patients WT for *ERBB2*. TNR-based SCNA signal is indistinguishable from negative controls at 0.1% dilution. (B) Receiver operating characteristic (ROC) curves for `dryclean` and TNR demonstrate increased sensitivity *in silico* dilutions of tumor DNA from *ERBB2* amplified breast cancers, with highest improvement and sensitivity in WGS. (C) *in silico* dilution results from 34 randomly selected GBM WGS samples for which SCNA signal was aggregated across 34 loci known to be recurrently amplified in GBM. These results show a similar improvement in tumor DNA detection for `dryclean` relative to other approaches for as low as 0.1% dilution.

Specifically, `dryclean` outperformed TNR across platforms for dilutions as low as 1 in 1000 in WGS. This suggests that `dryclean` (in combination with a primary tumor profile and a suitable PON) can significantly improve the detection of microscopic burdens of tumor cells and clinically relevant SCNAs in relapse tissue.

To further explore the utility of `dryclean` in detecting ctDNA in the absence of a primary tumor profile, we carried out *in silico* dilution experiments on 34 GBM WGS samples from TCGA. For tumor detection, we chose a panel of previously described (32) frequently amplified genes in GBM as markers across which we aggregated `dryclean` and TNR signal. We found that `dryclean` was able to reliably detect the presence of tumor at as low as 1% dilution (area under curve, AUC = 0.90), greatly exceeding the performance of TNR (AUC = 0.69) (Figure 6C). These results suggests that `dryclean` enrichment of foreground SCNA signal can improve tumor detection even in the absence of a primary tumor WGS profile.

## Discussion

We have demonstrated that `dryclean` improves the accuracy of SCNA detection across multiple sequencing platforms. However, we note that `dryclean` is not a method for SCNA characterization, but an algorithm for read depth normalization. In this study, we combine `dryclean` with CBS (CBS-DC) (24), primarily due to ease of implementation. Additional performance gains over CBS-DC might be obtained by combining `dryclean` with other, more sophisticated, SCNA detection algorithms (e.g. TITAN, FACETS) (13, 14) through minor bioinformatic modifications that replace the TNR input of these methods with the `dryclean` foreground signal.

The results obtained from `dryclean` greatly depend on the composition of the PON, which ideally should reflect the spectrum of assay and batch specific bias that is observed in a new tumor sample. Increasing the number of normal samples in the PON will incrementally improve the quality of results (Supplementary Figure S6, see Methods), but larger PONs require more compute to train (in the batch learning step (Figure 1)). Given standard computational limitations (storage, RAM, CPU), it is useful in practice to build a PON of 50-500 samples that is balanced with respect to common noise modes. If the PON is not representative of the assay and batch patterns in the tumor samples being analyzed, `dryclean` will produce sub-optimal results. This is especially true if the PON and tumor samples are derived from different sequencing platforms (e.g. HiSeq 2500 vs. HiSeq 4000 vs. NovaSeq). A notable use case of `dryclean` is in the analysis of cancer cell lines for which a matched normal can’t be found but for which a reasonable PON exists. We did not explore applying `dryclean` to the analysis of germline CN in this study; however, we believe that it may prove similarly useful in the sensitive detection of rare constitutional CN variants in clinical samples and populations.

Our `dryclean` analyses of cancer WGS implicate replication timing in the commonly noted “wave artifacts” in cancer sequencing data. Our analyses raise the possibility of using `dryclean` to chart the landscape of replication timing in cancer samples, including the nomination of somatic replication timing quantitative trait loci (21). However, since the `dryclean` background is inferred from a panel of normals, a replication timing-inference pipeline that depends exclusively on this signal may miss important replication timing alterations that arise in lineages that are not represented in the PON or result from large-scale somatic rearrangements. We foresee that additional processing downstream of `dryclean` foreground and background signals will be required to derive an optimally accurate replication timing signal from cancer WGS.

Our WGS dilution results suggest that `dryclean` may help dramatically increase the sensitivity of SCNA detection in the setting of cancer screening or relapse monitoring, including solid biopsy as well as liquid biopsy of cell free plasma or peripheral blood (e.g. clonal hematopoiesis) (5, 7, 31). Since previously published ctDNA methods can only reliably detect tumor-specific variants at an allelic or tumor-cell fraction > 0.1 (WES, WGS), our results suggest that `dryclean` may increase the sensitivity of ctDNA tumor detection as much as 10-fold over state-of-the-art approaches (6, 31). Since our *in silico* dilutions were performed on solid tissue-derived sequencing data, additional studies will be required to validate the utility of `dryclean` on liquid biopsy. This will include the generation and application of a ctDNA PON that captures the unique library characteristics of this assay (e.g. fragmentation patterns (33)).

Though we have demonstrated the use of `dryclean` on WES, TP, and WGS data from TCGA, we believe `dryclean` can improve the detection of SCNA across a variety of sequencing platforms, not just standard DNA sequencing. Future applications of `dryclean` may also include single-cell assays (e.g. scCNV, scATAC) that produce thousands of genomic profiles in a single experiment but may suffer from significant read depth bias (e.g. whole genome amplification, transposase accessibility) as well as sparsity (34). Application of `dryclean` in these data may help to better characterize genomic heterogeneity and link single cell SCNA patterns to phenotypic readouts of cell state.

## Conclusion

Read depth bias limits the accuracy of SCNA detection in clinical and research cancer DNA sequencing. In this study, we have introduced and benchmarked `dryclean`, a method to remove assay– and batch-specific bias from read depth data across a several sequencing platforms (WES, TP, WGS). These results indicate that `dryclean` could be a powerful addition to TP, WES, and/or WGS clinical oncology bioinformatics pipelines. Analysis of `dryclean`-derived background indicate that biological (i.e. replication timing), rather than technical factors (i.e. GC bias during library preparation) drive the “wave artifact” commonly observed in cancer WGS. Analysis of *in silico* tumor dilutions simulating solid or liquid biopsy sequencing data indicates a potential role for `dryclean` in sensitive relapse detection.

## Methods

### Mathematical formulation of dryclean

A basic premise of `dryclean` is that assay and batch-specific bias define a low-rank subspace in high-dimensional read depth data, which we represent as a matrix of *m* genomic bins across *n* samples. We achieve this through the application of robust principal component analysis (rPCA), which can accurately separate low-rank structures and high-amplitude outliers in complex multivariate data (18). A natural application for rPCA is motion detection in video data, where each data vector represents a video frame of a figure moving against a mostly stable background.

rPCA decomposes an input matrix 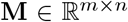 into a low-rank matrix 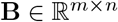 and sparse matrix 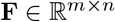 through the solution of a constrained optimization described in Eq. 1. This can be solved using iterative optimization methods such as Accelerated Proximal Gradient (APG) or Augmented Lagrangian Multiplier (ALM). Details of these algorithms can be found in (35, 36) respectively.

In general, these approaches minimize the objective function:

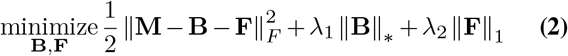

where λ_1_, λ_2_ are tuning parameters set to 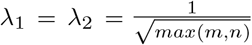, and 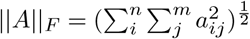 is the Frobenius norm of matrix **A**.

The robustness of rPCA comes at some computational cost. Training of Eq. 1 requires in-memory access to the full data matrix at every iteration of the optimization. This limits performance on larger data (35), e.g. read depth matrices comprising millions of bins and thousands of samples. This “batch” inference is also inefficient for the analysis of new data (e.g. a new tumor sample), which requires the model to be repeatedly refit to previously analyzed data points. To address this issue, we employ the online rPCA (OR-PCA) approach introduced by Feng et al (37), which allows efficient inference of additional data vectors, given an initially trained matrix.

### dryclean pipeline

We combine the batch and online rPCA approaches in the `dryclean` pipeline (Figure 1). The first step in the pipeline is to infer a low-rank subspace from a PON read depth data. Let 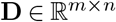 denote a matrix of log transformed read depth data across *n* samples and *m* bins, where each bin represents an exon, amplicon, hybrid capture probe, or tiled partition of the genome.

Applying the rPCA decomposition in Eq. 1 to **D**, we infer a low-rank matrix 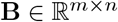 and a sparse matrix 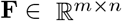. **B** is a low-rank representation of the PON, which comprises assay– and batch-specific noise (e.g. hybridization probe bias) but also can include common germline copy number polymorphisms (CNPs). The **F** matrix represents germline CN variants that are private to a few samples.

In practice, for inference, we use an online implementation that approximates the solution to (Eq. 1) for each tumor by projecting the tumor onto the subspace spanned by the *n* columns of the **B** and **F** matrix computed for just the PON. The latter then only needs to be computed once during a more (time and computationally intensive) training step.

`dryclean` is executed in three steps: First, read counts from a sample `.bam` are binned and corrected for GC content and mappability biases using a 2-step loess regression, in an implementation similar to HMMcopy (17). In the second step, a user defined number of normal samples are used to create a PON and extract the **B** matrix. Finally, using the **B** matrix, SCNA signals (foreground) for each tumor are extracted and stored as a Genomic Ranges object. Optionally, `dryclean` will use all normal samples to identify markers that commonly exhibit germline variation in a given dataset (Figure 1).

Though `dryclean` is a method for normalization of high-density read depth data, we also provide segmented output through the application of circular binary segmentation (CBS) to the **f** vector. We use these `dryclean`-derived SCNA calls throughout the manuscript to compare the quality of `dryclean` results to state-of-the-art methods; however, we foresee that `dryclean` output could be fed to more sophisticated segmentation algorithms, e.g. those employing additional data modalities such as heterozygous SNP counts or structural variant junctions to map SCNA with even higher accuracy and sensitivity. The contents of **f** vector represent the “foreground” of read depth data and captures the SCNAs while the **b** vector captures biases such as batch effect assay biases and replication timing which we term as “background”. As is shown in subsequent sections both vectors provide insight into different aspects of tumor biology.

### PON optimization

`dryclean`’s performance depends on the choice and number of normal (i.e. euploid) samples used to create the **B** matrix (PON). To objectively assess this performance, we use a measure of read depth noise called derivative log ratio spread (DLRS). It is a standard measure of the quality of log ratio (logR) data in SCNA analysis. It gauges marker to marker consistency in the signal and a larger spread is indicative of poor quality logR data (38). Applying DLRS across our benchmark analyses, we see that noise decreases as the PON size increases. Notably, we see a significant reduction in noise compared to TNR with a PON composed of as few as 50 samples (Supplementary Figure S6).

`dryclean` provides two options for sample selection for users wishing to generate smaller PONs from a larger dataset of normal samples. The first option is to build a PON from a randomly selected set of normal samples. A drawback of this approach is that less frequent batch effect modes may not be represented in the PON and thus `dryclean` will fail to correct coverage in tumor samples affected by those modes. Though one can build a very large and representative PON, the computational cost in building and applying such a dataset (even in batch rPCA mode) may be prohibitive. To address this issue, we provide users with a second option to create a PON that is balanced across batch effect modes. This is achieved efficiently by clustering on the basis of the low-rank signal (**b** vectors) computed for an rPCA decomposition performed across a small subset of genomic regions and a full set of normal samples and clustering these vectors. The normal samples to be used in the PON are then selected so that each cluster is represented. We see that there is rich structure among the normal samples (Supplementary Figure S7). This allows users to keep the size of the PON as small as possible while maximizing the information in the PON.

For all WES and TP `dryclean` analyses reported in the manuscript, we used a PON of 100 samples selected by the hierarchical clustering approach. To assess the performance of `dryclean` on WGS data (CCLE and TCGA), we generated a PON of 400 normal samples from the TCGA WGS cohort using the clustered approach

### CNV filter

Since `dryclean` uses a PON and not individual paired normal samples, an additional step is required to separate the SCNA signal from common germline CN polymorphisms (CNP’s) or matched normal germline CN variants (CNV’s). `dryclean` performs this step after creating the PON. Each normal sample is decomposed against the PON and the **f** vectors are extracted. This vector captures the germline CNV / CNP as opposed the **b** vector that encodes the background. Supplementary Figure S8 shows a small region of the genome with all these vectors concatenated in column space. `dryclean` creates an **f** vectors for all normal samples and then, using user defined frequency thresholds, annotates the markers harboring germline CNV and CNP, which are removed from downstream SCNA recurrence (i.e. GISTIC 2.0 (39)) analysis. Supplementary Figure S9 shows the reduction in narrow peaks from GISTIC 2.0 after this filter is applied.

### Data sources

Tumor and normal Illumina Hi-Seq `.bam` files were obtained from The Cancer Genome Atlas (TCGA, dbGAP identifier: phs000178.v1.p1) (40) via the GDC portal (https://portal.gdc.cancer.gov/). For WGS analyses, we used 943 samples with 23 different cancer types. For WES analysis, we used 1576 randomly selected samples comprising 19 different cancer types. We simulated targeted panel (TP) sequencing in tumor and normal samples by sub-sampling “target regions” from TCGA WES coverage data. We used 572 Cancer Gene Consensus (41) genes as targets. The distribution of the samples for different platforms can be found in (Supplementary Figure S10). 320 cancer cell line encyclopedia (CCLE) WGS were obtained from the CCLE portal (www.broadinstitute.org/ccle).

### Benchmarking

We compared circular binary segmentation (CBS) (24) ran on `dryclean` (CBS-DC) against two commonly used approaches for SCNA calling, CBS ran on TNR (CBS-TNR), and FACETS (14). For CBS-TNR, we used log-transformed, GC and mappability corrected TNR as an input. For CBS-TNR in CCLE cell lines that lack a matched normal, we computed a “pseudo-TNR” whose denominator was a “composite diploid sample” obtained by averaging GC and mappability-corrected read depth profiles across 943 TCGA WGS normal samples.

FACETS employs allele-specific copy number analysis of reference and variant allele read counts for germline polymorphic markers annotated by dbSNP (200K markers for WES and 7,400 markers for simulated TP). To assess the quality of WES and TP analyses, we used results from Affymetrix Genome-Wide Human SNP Array 6.0 (SNP6). SNP6 and other high-resolution microarray technologies are routinely used in basic research and clinical practice to efficiently detect SCNAs and represent the gold standard in clinical cytogenetics labs (26).

For all copy number comparisons, we removed centromeric segments, which are not reliably called by any SCNA method, including SNP6. We also limited our analysis to segments of 1MB or larger in size, since the reliability of more focal SCNA calls on any platform (including SNP6) is imperfect (14). For segment concordance, we defined concordant segments as those with at least 70% reciprocal overlap between both the start and end positions in the subject and query. To assess CN concordance, we used standard thresholds on segment intensities (i.e. segment mean value for CBS-DC and CBS-TNR or “cnlr.median” field in FACETS) to assign segments a “CN status”, i.e. gained vs lost. Specifically, we classified segments with intensities > 0.1 as “gains” and < −0.1 as “losses”. We then measured the proportion of correct “gain” vs. “loss” designations across all genomic coordinate-concordant segments (from above).

### ctDNA WGS simulations

For breast cancer (BRCA) samples, we simulated WGS dilutions at 5 dilution levels (*α* = 10%, 1%, 0.1%, 0.01% and 0%), 10 replicates, and 5 cases for a total of 250 samples. For each sample, dilution *α*, and 1 Kbp bin *i* with tumor and peripheral blood read count *t_i_* and *p_i_*, we sampled spiked tumor counts 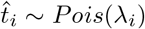 from a Poisson distribution with 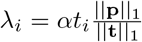. The simulated dilution read counts *s_i_* for each bin *i* were then defined as 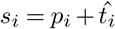. We then processed read counts through `dryclean` or TNR and analyzed read density at the *ERBB2* locus across various dilutions. For each simulation, we used a randomly chosen blood sample. Similar to BRCA, for glioblastoma (GBM) we used 34 samples and simulated dilutions at 5 dilution levels (*α* = 10%, 1%, 0.1%, 0.01% and 0%).

### Replication timing analysis

We ran `dryclean` on 320 CCLE WGS samples. For each sample, the inferred background (**b** vector) was then collapsed at 1 Mb level and median smoothed. We calculated the replication “score” by taking the inner product of **b** vector with the lymphoblastoid cell line (LCL) replication timing profile (21), an average of 6 replication timing profiles derived from 1000 genomes LCLs. We then classified cell lines as high (score greater than 75th percentile), moderate (score between 75th and 25th percentile) and low replicating (score less than 25th percentile). Finally, we validated this classification by comparing the expression of genes associated with cell proliferation (23) within these groups.

### Availability of data and software

`dryclean` is available in the GitHub repository https://github.com/mskilab/dryclean Data used to generate these results are based upon data generated by the TCGA Research Network: https://www.cancer.gov/tcga. (TCGA, db-GAP identifier: phs000178.v1.p1). All the CCLE processed datasets are available at the CCLE portal (www.broadinstitute.org/ccle) and DepMap portal (http://www.depmap.org). Raw sequencing data are available at Sequence Read Archive (SRA) under accession number PRJNA523380.

## Supplementary File

### Supplementary Figures

**Supplementary Figure S1.**
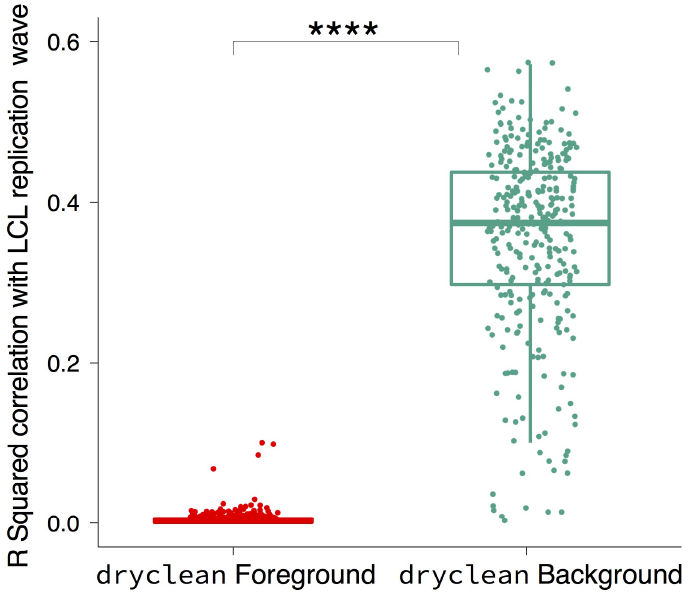
Correlation of dryclean with LCL replication wave. Higher correlation of `dryclean` background with LCL which is much higher than `dryclean` foreground that comprises of SCNAs. Each point here represents a CCLE cell line

**Supplementary Figure S2.**
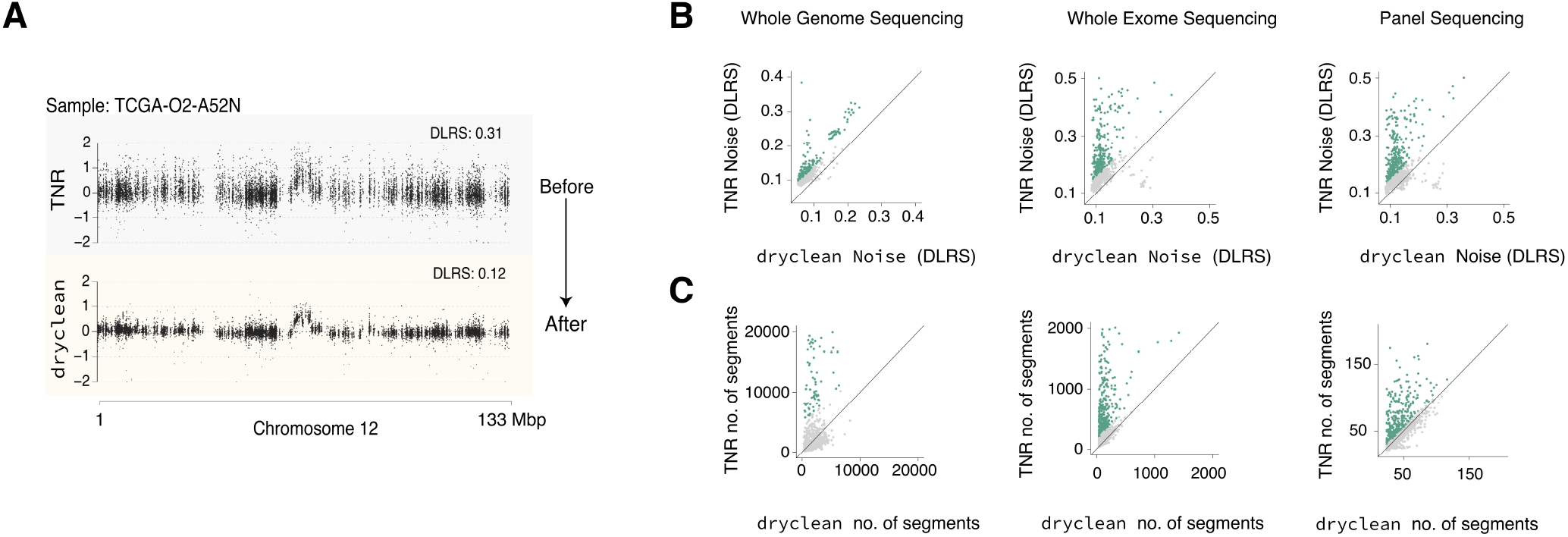
dryclean reduces read depth noise in WES and TP. (A) A representative WES sample with decreased DLRS in `dryclean` (yellow panel) relative to TNR (green panel). The higher DLRS in the TNR indicates a high burden of assay and batch specific noise. (B) Pan-cancer comparison of DLRS between `dryclean` and TNR in WES and simulated TP across 1576 TCGA cases. Each data point represents a TCGA case. For both platforms, DLRS is significantly reduced by `dryclean` relative to TNR. (C) Pan-cancer comparison of hypersegmentation (number of CBS segments) shows a reduction in `dryclean` results relative to TNR for both WES and TP. Each data point represents a TCGA case.

**Supplementary Figure S3.**
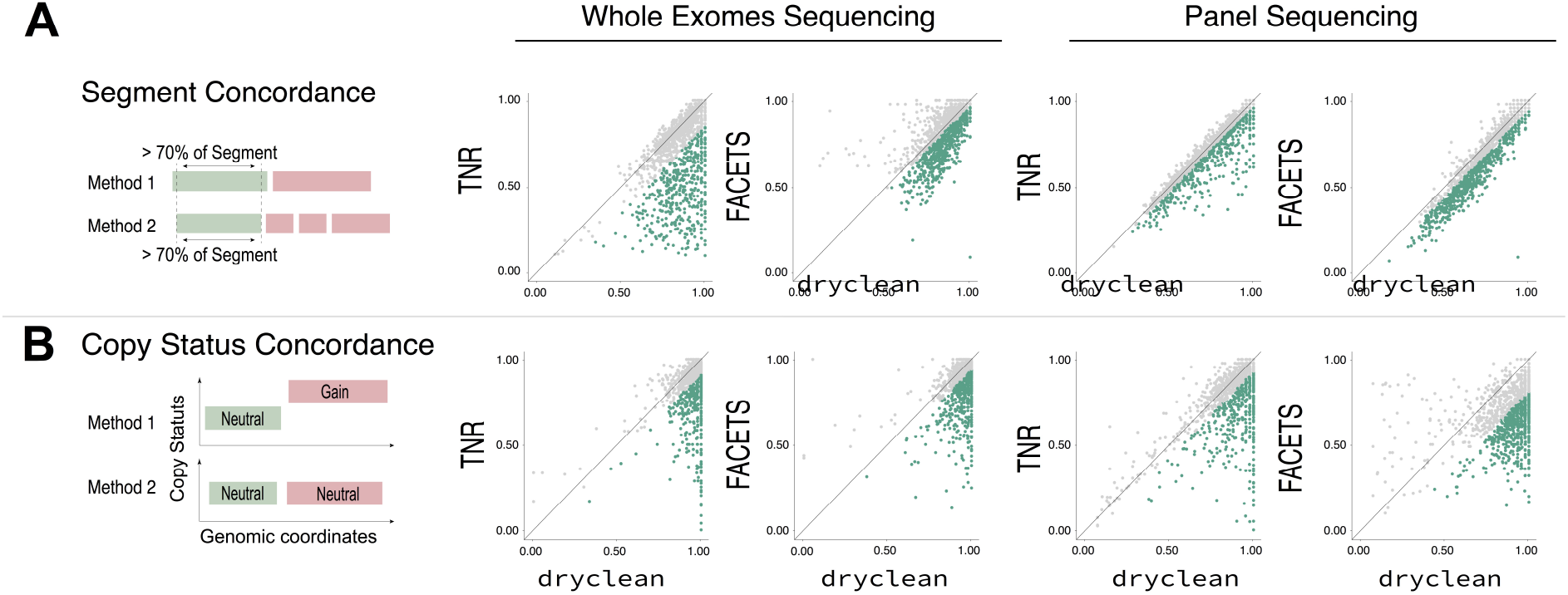
Concordance with SNP6. (A) Proportion of concordant segments with respect to genomic position. `dryclean` shows higher number of concordant segments as compared to FACETS and TNR in WES and to a lower extent in TP. (B) proportion of concordant segments with respect to copy number signal is much higher in `dryclean` in both WES and TP.

**Supplementary Figure S4.**
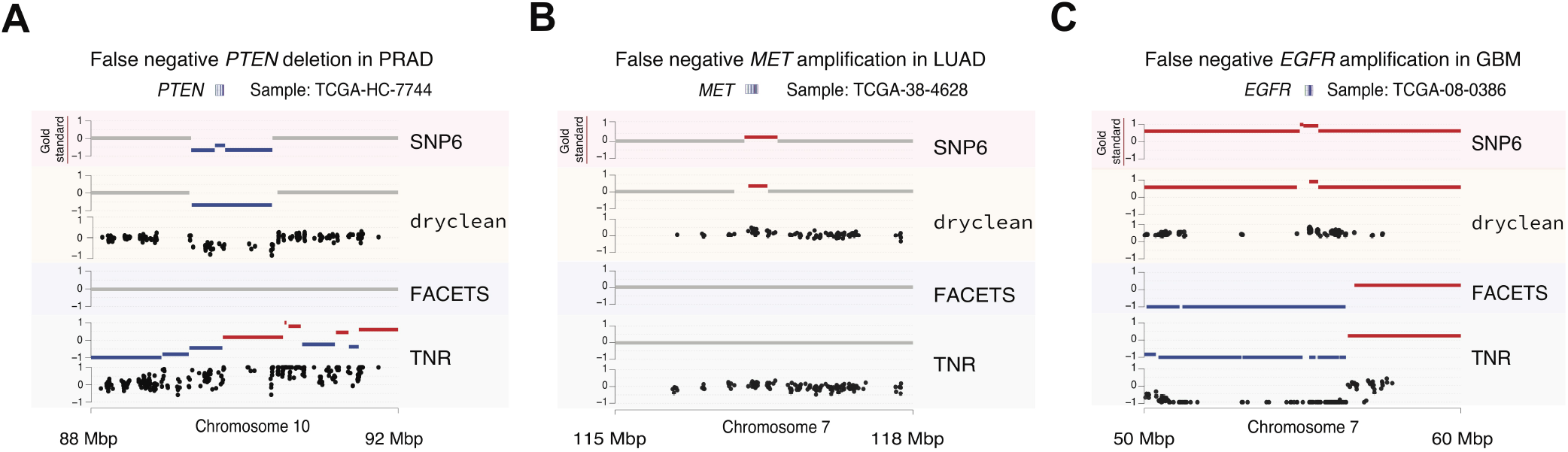
False negative examples of clinically important SCNA. (A-C) show instances of `dryclean` rescue of FACETS and/or TNR false negative calls in clinically actionable genes in WES data. (A) A SNP6 verified *PTEN* deletion is missed by TNR and FACETS due to noisy read depth signal, but caught by `dryclean`. (B) *MET* amplification is missed by TNR and FACETS but caught by `dryclean`. (C) A missed *EGFR* amplification in FACETS and TNR due to an artifactual event in the normal sample which skews the TNR. Both `dryclean` and SNP6 detect this event. For `dryclean` and TNR both raw data and segmentations from CBS are shown.

**Supplementary Figure S5.**
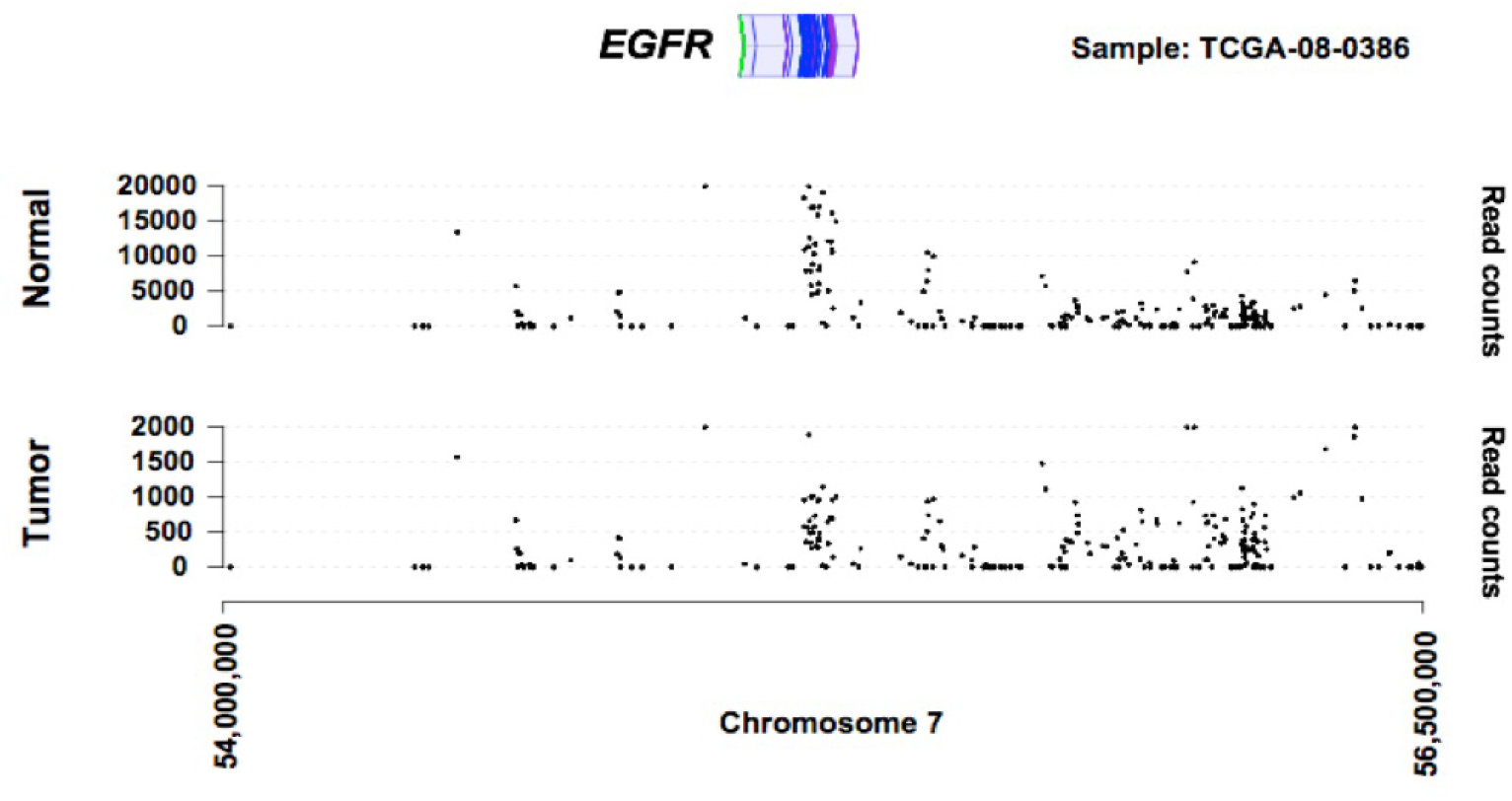
Paired normal sample shows a broad and high amplitude EGFR amplification. The two tracks show the raw read counts for tumor (lower track) and paired normal sample (upper track). It is clear from the read counts for these samples that the putative normal sample has 10 times as many reads mapping to *EGFR* locus. This explains the deletion signal seen in TNR and FACETS (Figure S4) even though they use different types of information, both methods ultimately normalize with paired normal sample.

**Supplementary Figure S6.**
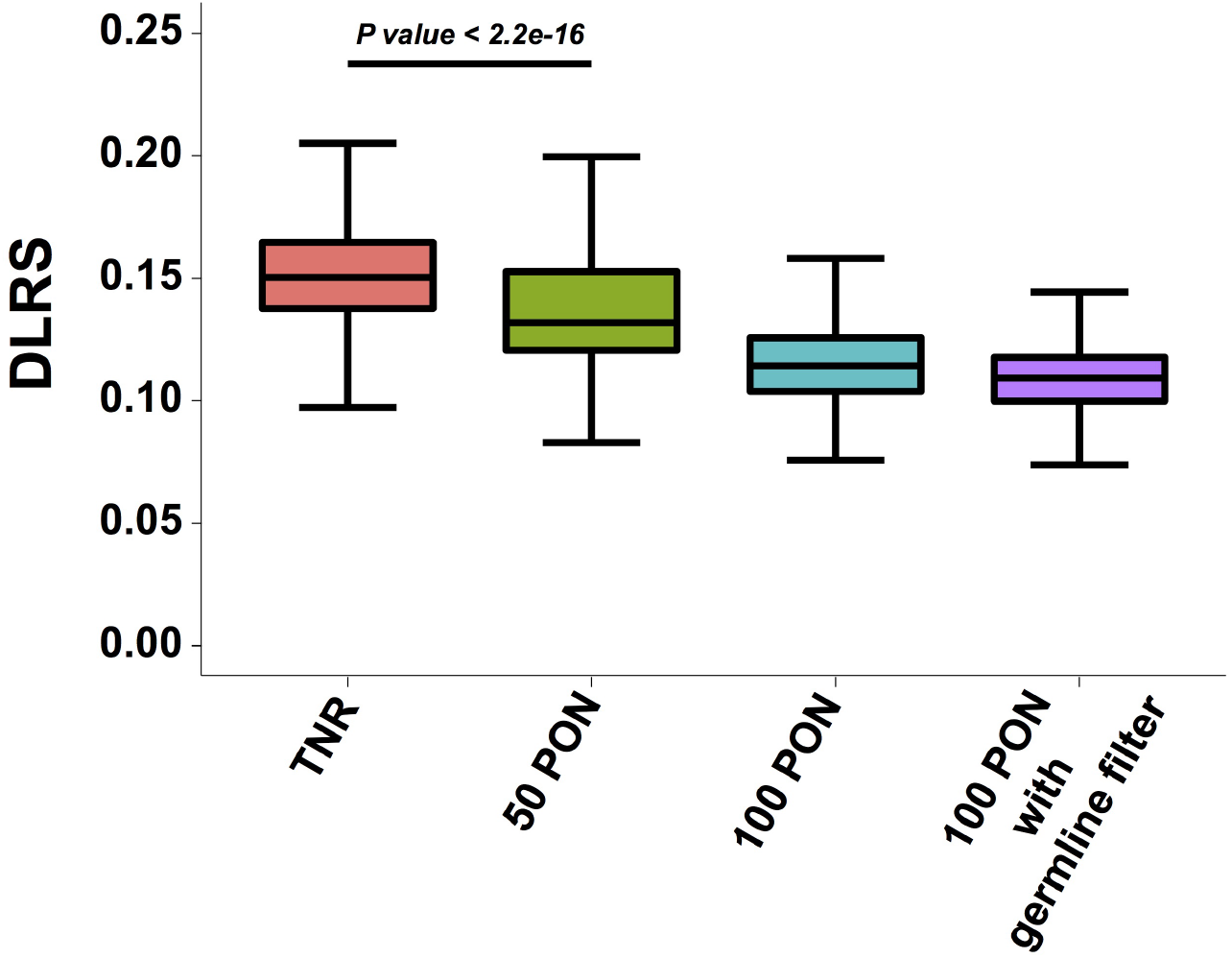
Quality of WES read depth data improves with number of normal samples in the PON. We mesure the quality of read depth data in terms of derivative log ratio spread (DLRS). We see reduction in DLRS value as the number of normal samples in the PON increase. Interestingly, there is a significant reduction in noise compared to TNR with a PON composed of as few as 50 samples (Wilcoxon rank sum test, p value < 2.2 × 10^−16^). When we add germline filter as described in main text, we get the best results. TNR: tumor-normal ratio, 50 PON: `dryclean` with 50 normal samples in panel of normal (PON). 100 PON: `dryclean` with 100 normal samples in PON.

**Supplementary Figure S7.**
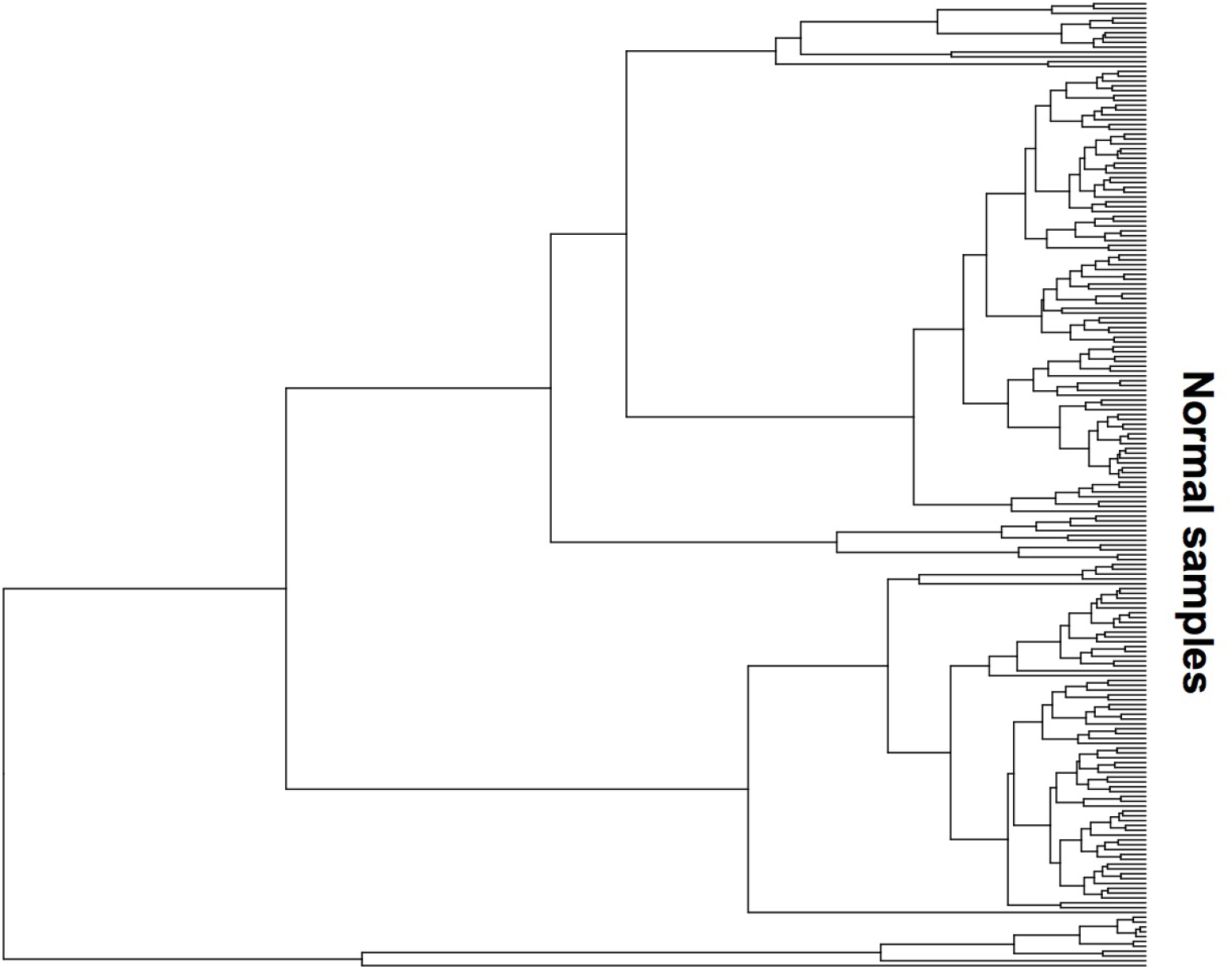
Rich structure in the genomic background subspace of WES normal samples. Shows the structure when 200 random WES samples were clustered using the inferred genomic background subspace. A small region of genome (chr. 22) was used to extract the background subspace. `dryclean` uses this approach to make a more informed decision to select normal samples to be included in PON.

**Supplementary Figure S8.**
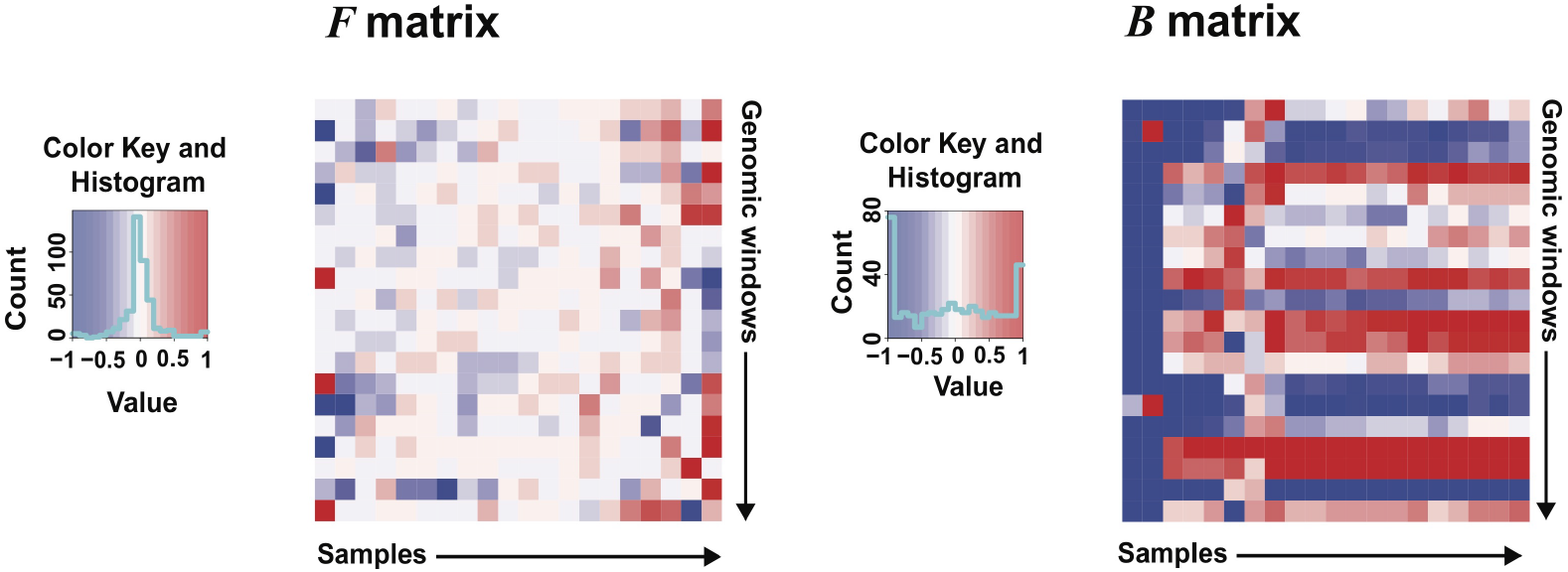
Visualization of B, F matrices after decomposition of PON. Each column here is a sample and each row is a genomic window. The values are normalized read counts in log space. The **F** matrix is mostly sparse and captures germline events. The **B** matrix captures genomic background. Here we see at least three distinct clusters, indicating the heterogeneity of biological and technical noise among samples.

**Supplementary Figure S9.**
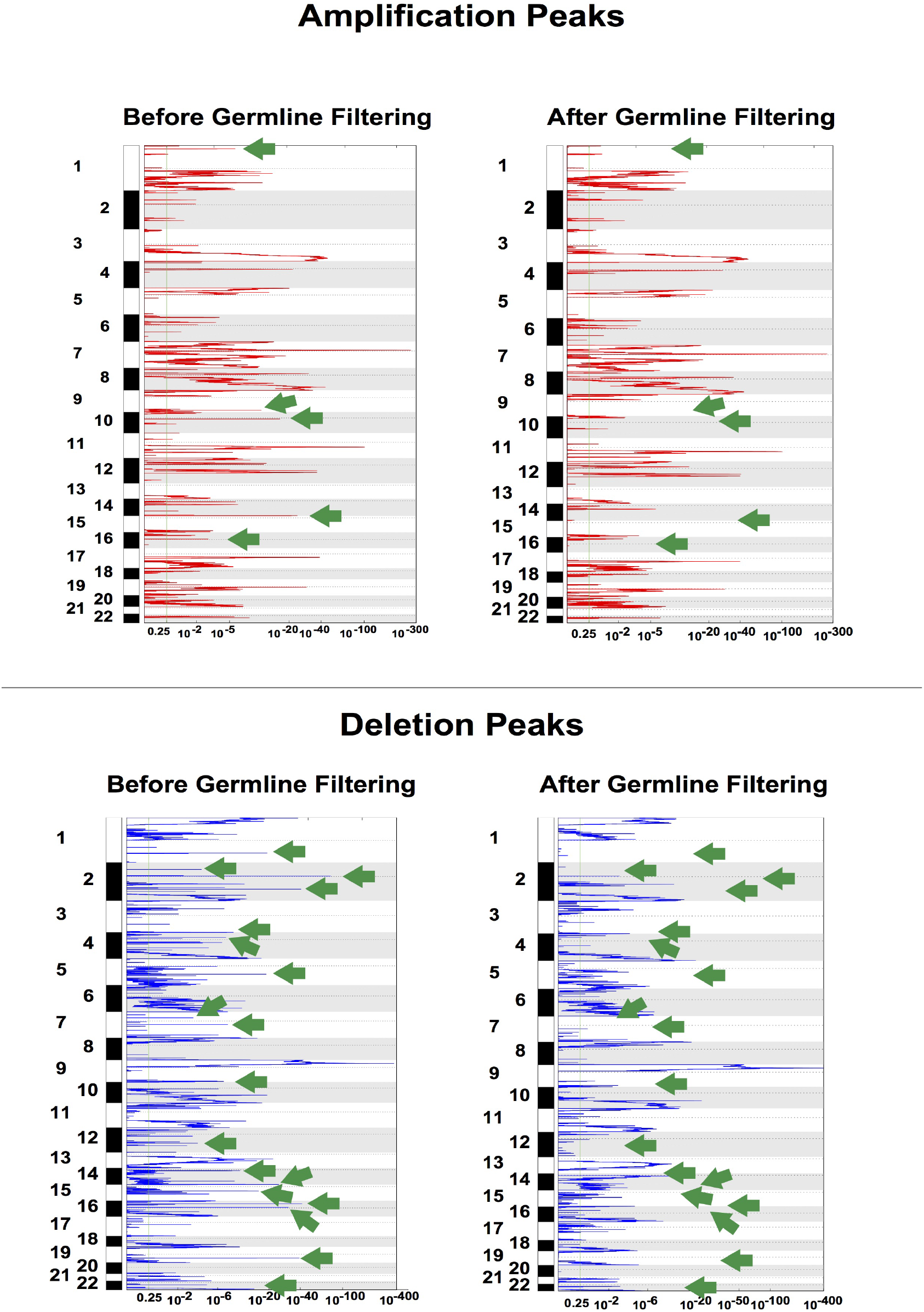
Effect of germline filter on the recurrent SCNA output from GISTIC2.0. The amplification and deletion peaks from `dryclean` before and after germline filter implementation within the algorithm. The green arrows point to the narrow peaks that are putative germline polymorphic events and are removed after application of the CNP filter.

**Supplementary Figure S10.**
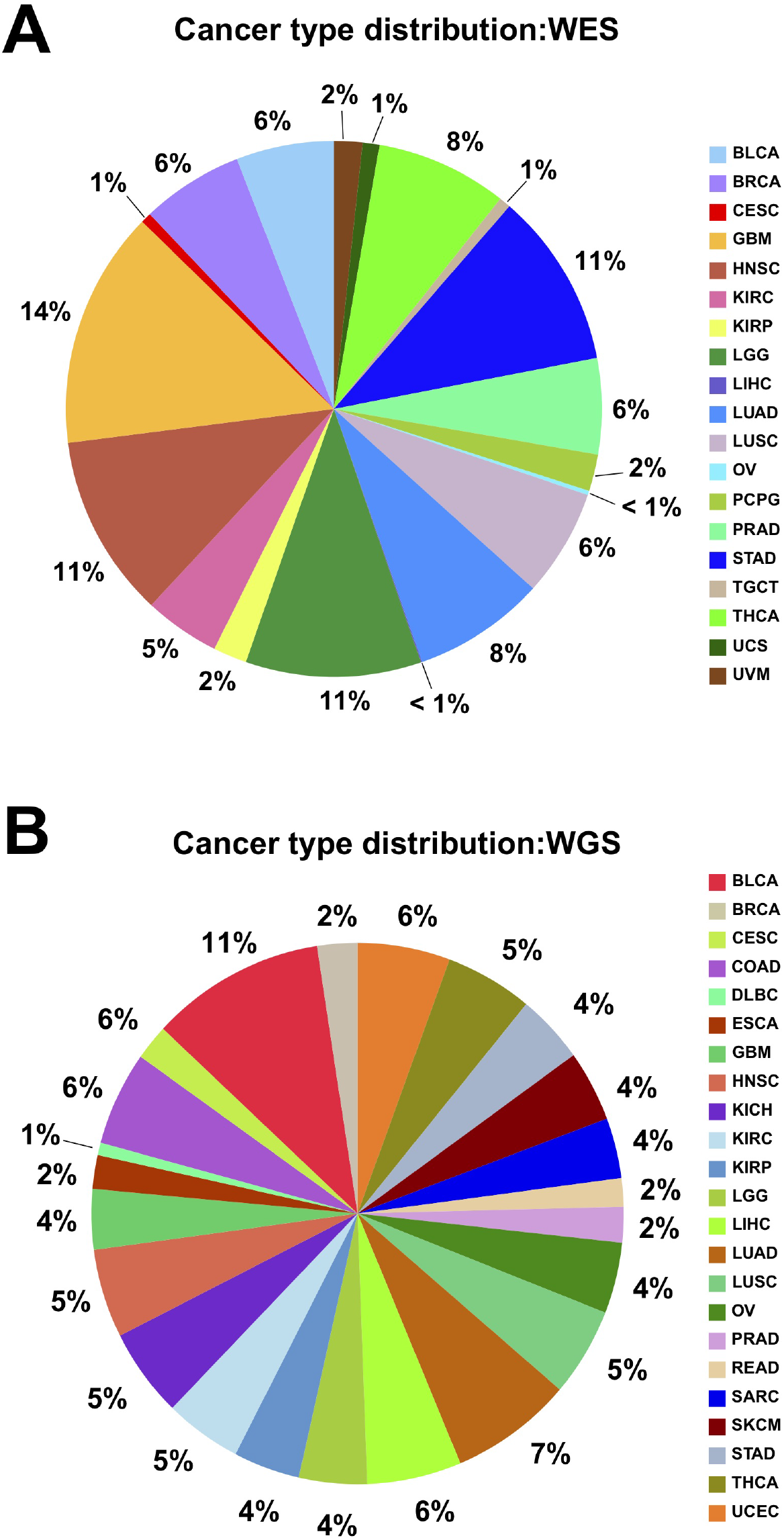
Summary of TCGA samples used in analysis. (A) Distribution of 19 tumor types in the WES sample set with 1576 samples. (B) Distribution of 23 tumor types in the WGS sample set with 943 samples

**Supplementary Table S1:**
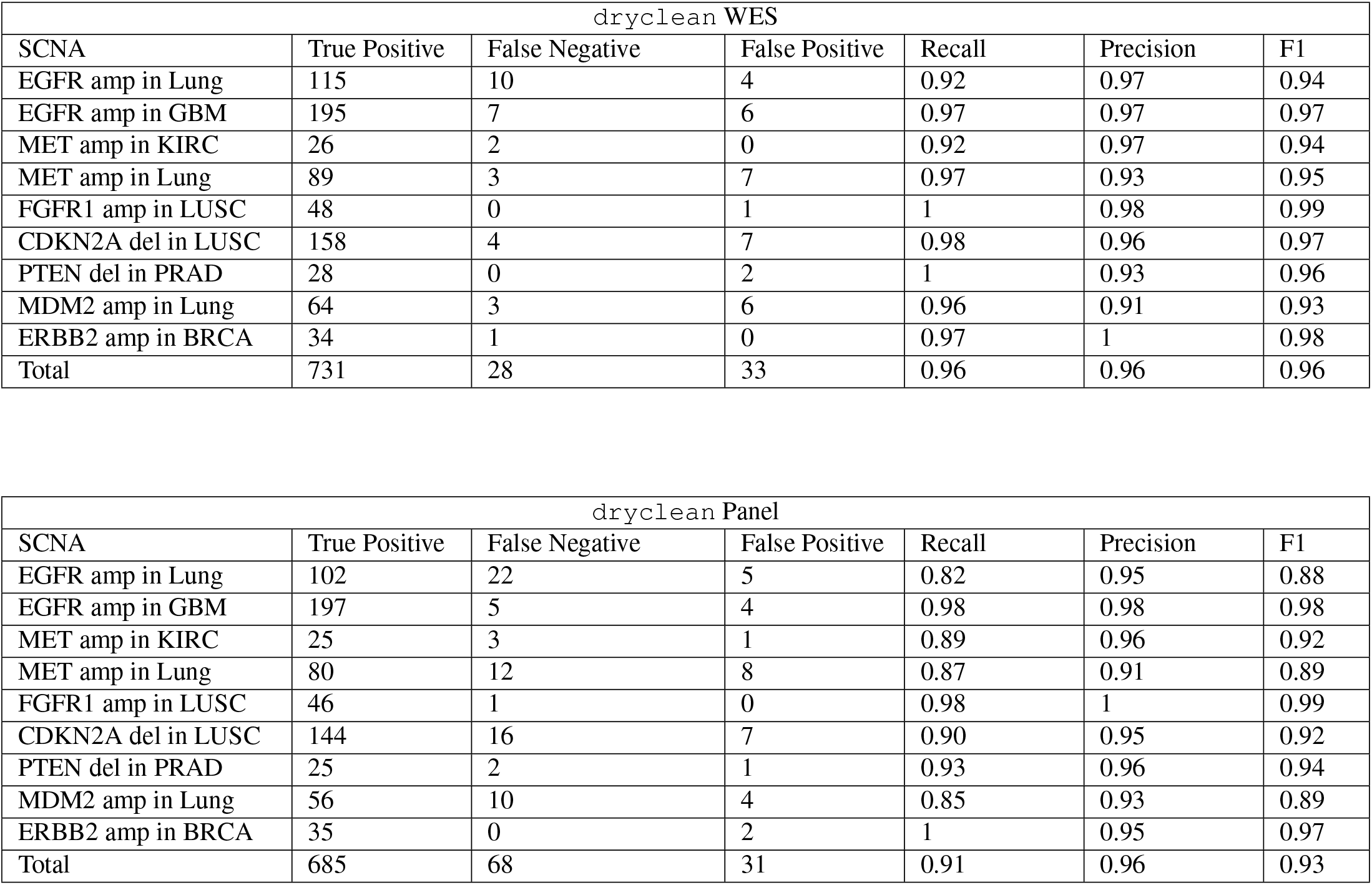
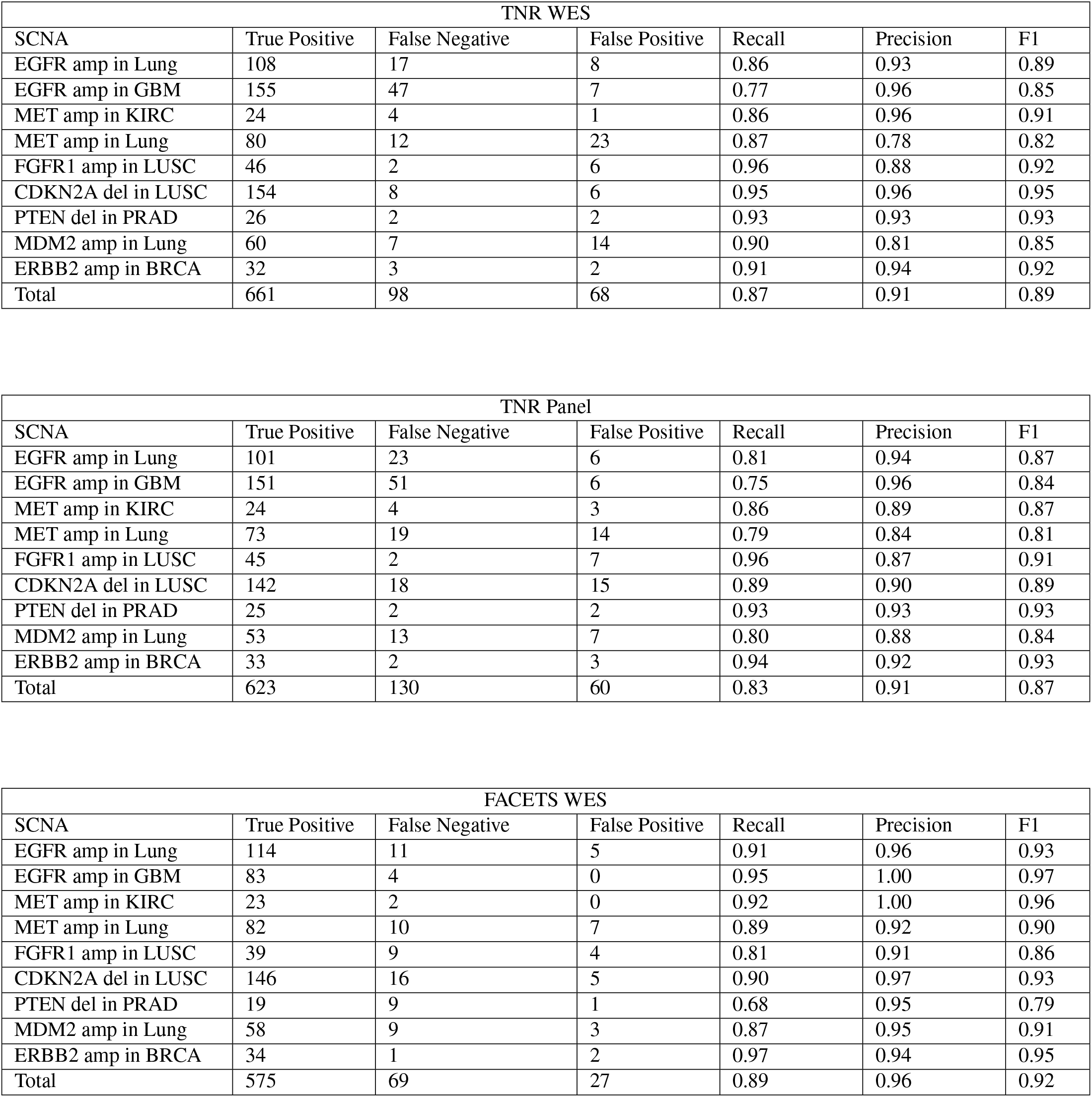
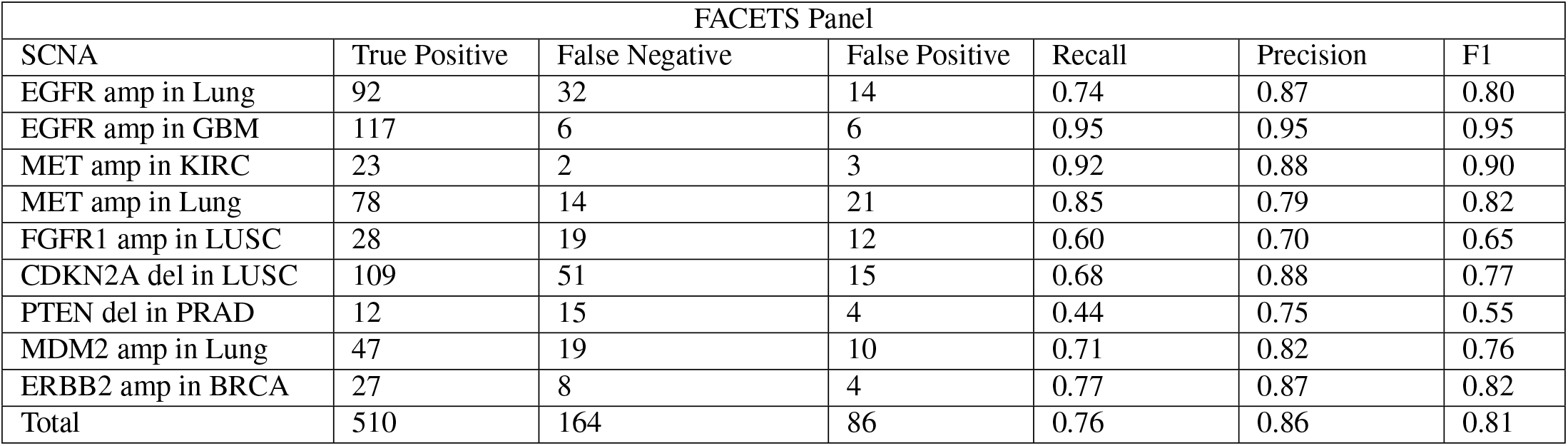
Raw counts for F1 calculations in Figure 5.

